# Visual recognition of social signals by a tecto-thalamic neural circuit

**DOI:** 10.1101/2021.08.17.456614

**Authors:** Johannes M. Kappel, Katja Slangewal, Dominique Förster, Inbal Shainer, Fabian Svara, Michal Januszewski, Shachar Sherman, Herwig Baier, Johannes Larsch

## Abstract

Social affiliation emerges from individual-level behavioral rules that are driven by conspecific signals^1–5^. Long-distance attraction and short-distance repulsion, for example, are rules that jointly set a preferred inter-animal distance in swarms^6–8^. However, little is known about their perceptual mechanisms and executive neuronal circuits^3^. Here we trace the neuronal response to self-like biological motion^9,10^ (BM), a visual trigger for affiliation in developing zebrafish^2,11^. Unbiased activity mapping and targeted volumetric two-photon calcium imaging revealed 19 activity hotspots distributed throughout the brain and clustered BM-tuned neurons in a multimodal, socially activated nucleus of the dorsal thalamus (DT). Individual DT neurons encode fish-like local acceleration but are insensitive to global or continuous motion. Electron microscopic reconstruction of DT neurons revealed synaptic input from the optic tectum (TeO/superior colliculus) and projections into nodes of the conserved social behavior network^12,13^. Chemogenetic ablation of the TeO selectively disrupted DT responses to BM and social attraction without affecting short-distance repulsion. Together, we discovered a tecto-thalamic pathway that drives a core network for social affiliation. Our findings provide an example of visual social processing, and dissociate neuronal control of attraction from repulsion during affiliation, thus revealing neural underpinnings of collective behavior.

## Main

Many animals live in groups, the result of a basic social affiliative drive that requires detection and approach of conspecifics. Social affiliation is a prerequisite of consummatory actions such as aggression, mating or play^3^, and is also a proximal cause of swarm, flock, and herd formation. While neuronal circuits mediating such behaviors have received much attention^3,14,15^, relatively little is known about the sensory detection of social signals (beyond pheromones)^15,16^, and how such cues feed into the regulation of social distance. One important class of visual social signals is biological motion (BM), which comprises conspecific movement patterns that trigger complex approach and pursuit behaviors^17–21^ and elicit a social percept in humans^9,10,22^. BM is also a key driver of zebrafish shoaling^2,11,23^, a collective behavior with well-characterized behavioral rules in groups or pairs of animals^2,6,11,24,25^, offering a model to investigate visual neural circuits underpinning social affiliation.

### Fish-like motion activates a conserved social behavior network

To identify the relevant neuronal circuits in 21 days old juvenile zebrafish, we generated unbiased maps of recent neuronal activity^13^ following shoaling with real or virtual conspecifics (Fig. 1a). Virtual conspecifics were projected black dots moving either with fish-like BM, or continuously, that are highly attractive and weakly attractive, respectively^2^ (Fig. 1b). After 45 minutes of shoaling, we recorded a snapshot of neuronal activity by rapid fixation and labeling of *c-fos (fosab)* mRNA^13^ using third generation *in situ* hybridization chain reaction^26^ (HCR) for volumetric fluorescence imaging through the entire depth of the midbrain, forebrain, and anterior hindbrain (Figs. 1a, c, S1). To compare neuronal activation across animals and conditions, we registered all image stacks to an averaged, age-matched, aldehyde-fixed brain and normalized the *c-fos* signal using a second HCR probe against pan-neuronally expressed *elavl3* (Fig. S1). Visual inspection of the merged *c-fos* signal from all animals identified 31 distinct clusters with robust activity in response to one or more stimulus conditions (Figs. 1c, S1).

**Fig. 1.**
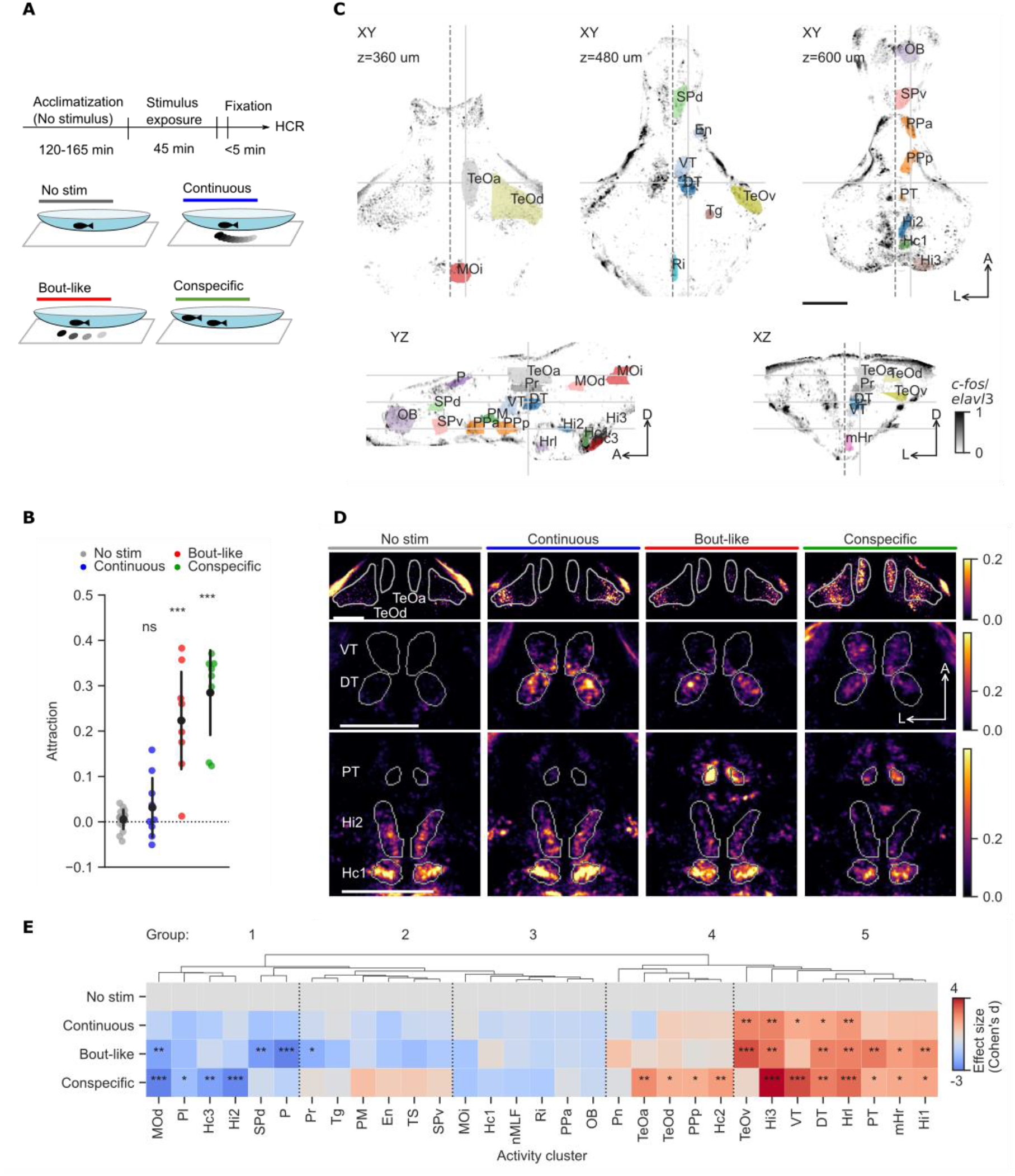
Fish-like motion activates a conserved social behavior network. **a,** Schematic of stimulus presentation before activity mapping: Individual animals interact with real conspecifics in a watch glass of 7 cm diameter or with a virtual conspecific consisting of a black dot projected onto a screen below each dish. **b,** Attraction is calculated as ((IADs-IADo)/IADs). IADo: observed inter animal distance, IADs: inter animal distance for time shuffled trajectories. n=8 animals (colored dots). Black dots represent mean±1SD. Asterisks represent significance levels of comparison to no-stimulus (No stim) group. NS: not significant, ***: p<0.001. Two-tailed t-test, Bonferroni corrected. **c,** Representative slices of maximum intensity normalized *c-fos* signal merged across all 28 registered animals. Views are horizontal (top row), sagittal (bottom left), and coronal (bottom right). Solid gray lines indicate corresponding planes across slices. Dashed line indicated midline. Scale bar: 200 μm. Arrows indicate A: anterior, L: left, D: dorsal. See also Fig. S1. **d,** Average normalized *c-fos* signal at three representative horizontal planes indicated in (C). n=6-8 animals per condition. Scale bars: 200 μm. **e,** Effect size of normalized bulk *c-fos* induction compared to no-stimulus condition was quantified as the mean difference divided by the pooled standard deviation (Cohen’s d). Negative values indicate lower *c-fos* signal than no-stimulus condition. Dendrogram represents hierarchical clustering. Asterisks indicate significance levels obtained from two-tailed t-tests in each activity cluster versus the no-stimulus group. p<0.05:*, p<0.01:**, p<0.001:***, Bonferroni-corrected per activity cluster. n=6-8 animals per condition. **Abbreviations: DT**: Dorsal thalamus; **En**: Entopeduncular nucleus; **Hc1**: Caudal hypothalamus 1; **Hc2**: Caudal hypothalamus 2; **Hc3**: Caudal hypothalamus 3; **Hi1**: Intermediate hypothalamus 1; **Hi2**: Intermediate hypothalamus 2; **Hi3**: Intermediate hypothalamus 3; **Hrl**: Rostral hypothalamus, lateral; **mHr**: Rostral hypothalamus, medial; **MOd**: Medulla oblongata, dorsal; **MOi**: Medulla oblongata, intermediate; **nMLF**: Nucleus of the medial longitudinal fasciculus; **OB**: Olfactory bulbs; **P**: Pallium; **Pl**: Pallium, lateral; **PM**: Magnocellular preoptic nucleus; **Pn**: Pineal; **PPa**: Anterior parvocellular preoptic nucleus; **PPp**: Posterior parvocellular preoptic nucleus; **Pr**: Pretectum; **PT**: Posterior tuberculum; **Ri**: Inferior Raphe; **SPd**: Subpallium, dorsal; **SPv**: Subpallium, ventral; **TeOa**: Tectum, anterior; **TeOd**: Tectum, dorsal; **TeOv**: Tectum, ventral; **Tg**: Lateral tegmentum; **TS**: Torus semicircularis; **VT**: Ventral thalamus.

Splitting the data by stimulus group revealed that social context differentially activated these clusters. Activation by real and virtual conspecifics overlapped in a subset of clusters including Hc1, Hrl, and Hi3 in the caudal, rostral and intermediate hypothalamus while showing a distinct pattern in other areas. In the optic tectum, virtual conspecifics activated a ventrolateral cluster (TeOv), matching the retinotopic representation of the ventrally projected black dot visual stimulus^27^. In contrast, real conspecifics activated the anterior and dorsal optic tectum (TeOa, TeOd) more strongly (Fig. 1d). Virtual conspecifics activated a cluster in the dorsal thalamus (DT). Real conspecifics additionally activated an anterior cluster in the ventral thalamus (VT) (Fig. 1d). The DT *c-fos* cluster overlapped with a gene expression hotspot of the gene *cortistatin* (*cort,* also known as *sst3*), which we co-labeled using a third, multiplexed HCR probe and used subsequently as a marker (Sherman et al., in preparation). While *cort* and *c-fos* were expressed in a similar number of DT cells (*c-fos*: between 38±20 and 150±20 cells for no stimulus and conspecific, respectively), *cort* (82±12 cells) expression was practically non-overlapping at the level of individual neurons (1.5±1.4 cells) (Figs. S2a, b). One cluster in the posterior tuberculum (PT) stood out as selectively active with bout-like motion and real conspecifics, the two conditions that elicited high social attraction. In contrast, *c-fos* in hypothalamic cluster Hi2 was highest for the no-stimulus condition and inversely related to stimulus attraction (Fig. 1d).

To quantify these trends, we calculated an average bulk *c-fos* signal per cluster in each animal (Figs. S2c, 1e). Statistical analysis of the bulk *c-fos* signal revealed significant modulation of activity in 19 clusters by at least one stimulus relative to the no-stimulus condition (Fig. 1e). Hierarchical clustering separated 5 major groups of clusters that were 1) suppressed by most stimuli, 2) weakly suppressed by virtual conspecifics, 3) not modulated relative to “no stimulus”, 4) activated more by real conspecifics than virtual ones, and 5) activated by most stimuli (Fig. 1e).

Together, this unbiased global activity map identifies a ‘social behavior network’ for shoaling whose activity is modulated by real and virtual conspecifics (group 5). Preoptic and other hypothalamic clusters in groups 1 and 4 that were not strongly modulated by virtual conspecifics suggest that real conspecifics elicit neuronal responses beyond those controlling acute social affiliation, including the perception of threat and homeostatic stress mechanisms, potentially via additional sensory modalities^13,16,28,29^. Thus, the set of clusters activated by virtual conspecifics highlights a core network to investigate the sensorimotor transformation of shoaling, beginning with the visual recognition of conspecifics.

### A cluster of neurons in the dorsal thalamus is tuned to fish-like motion

While our *c-fos* labeling method confirms a conserved contribution of hypothalamic components to the social behavior network^12,13^, it additionally highlights the visual pathway in TeO and DT through which fish-like motion signals enter the brain. To understand stimulus selectivity of individual neurons in these visual areas, we turned to volumetric two-photon calcium imaging of juvenile brain activity in response to presentation of virtual conspecifics.

Fish that expressed nuclear-localized GCaMP6s in almost all neurons (*elavl3:H2B-GCaMP6s*)^30^ were immobilized on the stage of a microscope equipped with a custom-built remote focusing setup for rapid image acquisition. We imaged simultaneously in 6 imaging planes at 5 Hz, extending 600 x 600 x 200 um (x,y,z) (Fig. 2a). This volume included the retinorecipient brain areas highlighted by our *c-fos* analysis, DT and TeO, and pretectum, nucleus isthmi, ventral thalamus and habenulae (Figs. 2b, S3b). All moving dot stimuli elicited widespread neuronal responses across the visual system. Analysis of registered responses across animals revealed that most responsive neurons reside in TeO (44%), pretectum (16%), DT (13%) and nucleus isthmi (11%) (Fig. S3), quantitatively matching the *c-fos* mapping results (Fig. 2c).

**Fig. 2.**
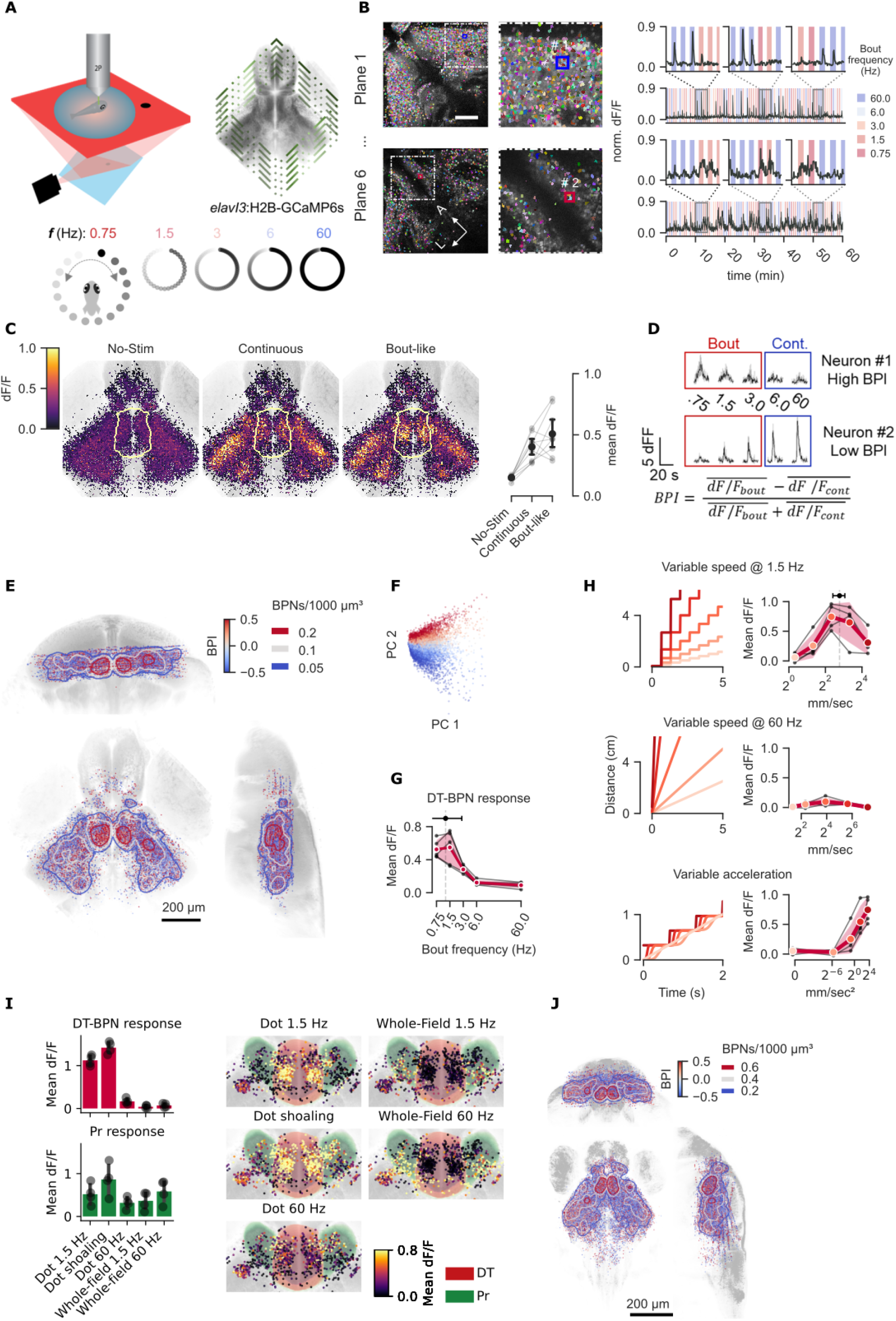
Dorsal thalamus neurons are activated by fish-like motion. **a,** Schematic of the 2-photon remote focusing experimental setup. Agarose-embedded animals saw dots in apparent motion at five bout frequencies. **b,** Example imaging planes in the tectum and dorsal thalamus with all segmented neuronal ROIs. Representative ΔF/F traces of one tectal and one thalamic neuron tuned to continuous (60 Hz) and bout-like (1.5 Hz) motion, respectively. Scale bar: 100 μm. **c,** Left: mean ΔF/F responses to no stimulus, continuous and bout-like motion of all recorded neurons (n=26057) from 9 fish (18-22 dpf) aligned to a juvenile average brain. Outline: Dorsal thalamus (DT). Right: Mean ΔF/F responses of all DT neurons per fish (n=306±217, 2756 total). **d,** Top: Mean ΔF/F responses of example neurons in **b** to all stimulus frequencies. Bottom: Bout preference index (BPI) is the normalized difference between responses to bout-like and continuous stimuli. **e,** Anatomical distribution of all BPNs colored by BPI. Contours are Gaussian Kernel Density Estimations (KDE) of all Bout Preference Neurons (BPNs, BPI > .5) indicating density of 0.05/0.1/0.2 BPNs / 1000 μm^3^. The highest density is located in DT. **f,** BPI distribution in tuning space defined by Principal Component Analysis (PCA). Colors indicate BPI as in **e**. **g,** Mean DT-BPN (n=100±60 cells per fish) tuning to stimulus frequencies from 0.75 to 60 Hz shown in red. N=5 animals. Mean of all neurons was 1.4 Hz ±1.7 Hz, n=498 neurons. Black lines represent means of individual animals. Data from subset of animals in **e** with number of recorded DT-BPNs>10. **h,** Mean DT-BPN tuning to average speed at 1.5 Hz or 60 Hz stimulus frequency, and tuning to acceleration at 1.5 Hz, respectively. Cartoons on the left illustrate stimulus displacement over time. Mean ± 1 SD of all neurons shown above (n=219 neurons). Black lines represent means of individual animals (n=4, fish, 55±19 neurons per fish). **i,** Left: Mean DT-BPN and PreT responses to local dot motion and whole-field motion. Circles show mean of individual animals (n=4 fish, 68±14 (DT-BPN), n=72±34 (PreT) neurons per fish). Right: Anatomical distribution of DT-BPNs and pretectal neurons. **j,** Anatomical distribution of BPI and Gaussian Kernel Density Estimation (KDE) of all Bout Preference Neurons (BPNs, BPI > .5) as in E for 7 dpf larvae.

To identify BM encoding neurons, we computed a bout preference index (BPI) as the normalized difference in the response to behaviorally attractive bout-like motion versus unattractive continuous motion (Fig. 2d). The vast majority of neurons did not differentiate between bout frequencies (mean BPI 0.03±0.1). However, 9±4% of all neurons scored BPI > 0.5, corresponding to a three-fold increase in ΔF/F for bout-like motion compared to continuous motion in these neurons. We focused our attention subsequently on these putative bout preference neurons (BPNs). Most recorded BPNs were located in TeO (35%) and DT (21%) (Fig. S3). We next computed a Gaussian kernel density estimation (KDE), that yielded DT as the anatomical area of highest BPN density (Fig. 2e). Within DT, BPNs were concentrated in a posterior cluster, overlapping with DT *c-fos* activity (Fig. 1c). In contrast, tectal BPNs were distributed broadly along the antero-posterior and dorso-ventral axes at lower density (Fig. S3). The similarity in *c-fos* and GCaMP signals suggests that virtual conspecifics in the open-loop configuration activate key circuits for social recognition even in immobilized animals. To test whether BPI captures a meaningful axis in tuning space, we performed principal component analysis (PCA) based on mean ΔF/F responses. BPI formed a gradient in PC space along the second axis, suggesting that the bout vs. continuous distinction is prevalent in the recorded neural population (14% variance explained, Figs. 2f, S3).

### DT-BPNs encode local swim bout-like acceleration

If DT-BPNs function as sensory drivers of shoaling behavior, their tuning should match specific parameters of BM. DT-BPNs had a response peak at a stimulus bout frequency of 1.4±1.7 Hz, closely matching the juvenile’s typical swim bout frequency of ∼1.25 Hz that most effectively triggers shoaling^2^ (Fig. 2g, S7c). To ask if BPNs encode acceleration or average speed of virtual conspecifics, we collected a separate dataset and systematically varied each parameter independently (Fig. 2h). At continuous motion, DT-BPNs were barely modulated by stimuli moving at 2 to 150 mm/s. At 1.5 Hz, DT-BPNs yielded maximal responses at 6.8±1.5 mm/s (Fig. 2h), similar to a juvenile’s typical swim speed at ∼5 mm/s and, again, matching the behavioral tuning^2^ (Fig. S7b). Morphing acceleration from continuous to bout-like along Gaussian speed profiles at fixed average speed of 5 mm/s and 1.5 Hz bout frequency modulated DT-BPN responses as a function of acceleration with a maximum at the highest possible acceleration of 12 m/s^2^ (projector limit). Taken together, DT-BPNs detect BM via periodic acceleration at fish-like speed and bout frequency, and are, thus, tailored for the detection of juvenile zebrafish during shoaling.

To relate DT-BPN responses to naturalistic visual percepts, we tested another set of animals with ‘dot shoaling’ stimuli on trajectories that recapitulate positions of conspecifics relative to a real focal animal during shoaling (Fig. S4a). DT-BPNs were strongly and persistently activated by such stimuli (Fig. S4c). Self-motion during shoaling also generates global motion with temporal dynamics similar to the fish-like cues. To ask if global motion activates DT-BPNs, we rotated whole-field stimuli with matched bout-like motion and spatial frequency (Fig. S4b). Global motion strongly activates pretectal neurons^31^ but not DT-BPNs themselves (Fig. 2i), suggesting the latter encode fish-like BM and not self motion-induced visual signals.

### Bout recognition is already established in the larval visual system

Zebrafish shoaling with real or virtual conspecifics emerges at around two weeks of age^2,6,24^, whereas younger fish show mainly inter-animal repulsion^2,32^. We therefore hypothesized that functional maturation of BPNs coincides with this transition. Contrary to this prediction, BPNs already existed in larvae, however with lower fractions compared to juveniles (6±2% of all recorded neurons). Further, BPNs were similarly distributed in the brain, with the KDE center in DT (Fig. 2j). Registration of the larval data to the Max Planck Zebrafish Brain (mapzebrain) atlas^33^ confirmed localization of the DT-BPN cluster to the *vglut2a-*positive DT area, ventrally touching the *gad1b* positive VT^34^. The larval DT-BPN cluster was molecularly defined by expression of *cort,* as seen in juveniles, and *pth2* (Fig. S4d), a gene whose thalamic expression tracks the density of conspecifics in zebrafish via mechanosensory signals^35^. Overlap with *pth2* raises the possibility of multimodal integration of conspecific signals in DT. Further, the mean frequency tuning curve across all DT-BPNs was similar to juveniles (Fig. S4e). Thus, functional BPN maturation precedes shoaling, and the developmental transition is either gradual in nature or requires a change downstream of BPNs. The presence of BPNs in pre-juvenile stages provides an opportunity to investigate the circuit with the experimental tools and resources available in larvae.

### Electron microscopic reconstruction reveals connectivity between BPNs, the tectum and the social behavior network

Across species, the thalamus acts as a gateway for state-dependent sensory information^34,36^. We hypothesized that DT-BPNs could serve that role for social cues, connecting visual brain areas and the conserved social behavior network^12^. To reveal the anatomy of the DT-BPN circuit, we analyzed an electron microscopic (EM) whole-brain dataset of a 5 dpf larval zebrafish, acquired at synaptic resolution (Svara et al., manuscript in preparation; see Methods). We registered the larval DT onto the EM volume to identify the cell body location of putative BPNs in the dorsal thalamus (Fig. S5a-c). We randomly selected and completely traced 34 cells in this region (32 in the left and 2 in the right brain hemisphere; Fig. 3a, b). All of these cells extended their primary neurite ventro-laterally and showed both dendritic and axonal arborizations in a thalamic neuropil region, posterior to retinal arborization field AF4 (Fig. 3b, c). In this region, we randomly selected presynaptic contact sites on putative BPN dendrites, and completely traced their partner neurons (Fig. 3c). Besides intra- and inter-thalamic connectivity between DT neurons (n=7; Figure S6a), we identified synaptic input from the ipsi- and contralateral nucleus isthmi (n=3; Fig. S6b) and from tectal periventricular projection neurons (PVPNs, n=7) (Fig. 3d). PVPNs send their axons ventrally through the postoptic commissure, and make ipsi- or contralateral connections to putative BPNs within the thalamic neuropil region. A single PVPN can be presynaptic to several putative BPNs (Fig. 3e), and a single BPN can receive input from several PVPNs, both ipsi- and contralaterally (Fig. S6c). Again, by tracing synaptic partners, we found that these PVPNs receive direct visual input from a specific class of retinal ganglion cells (RGCs), which all arborize in the SFGS3/4 layer of the tectum^37^ (Fig. 3f).

**Fig. 3.**
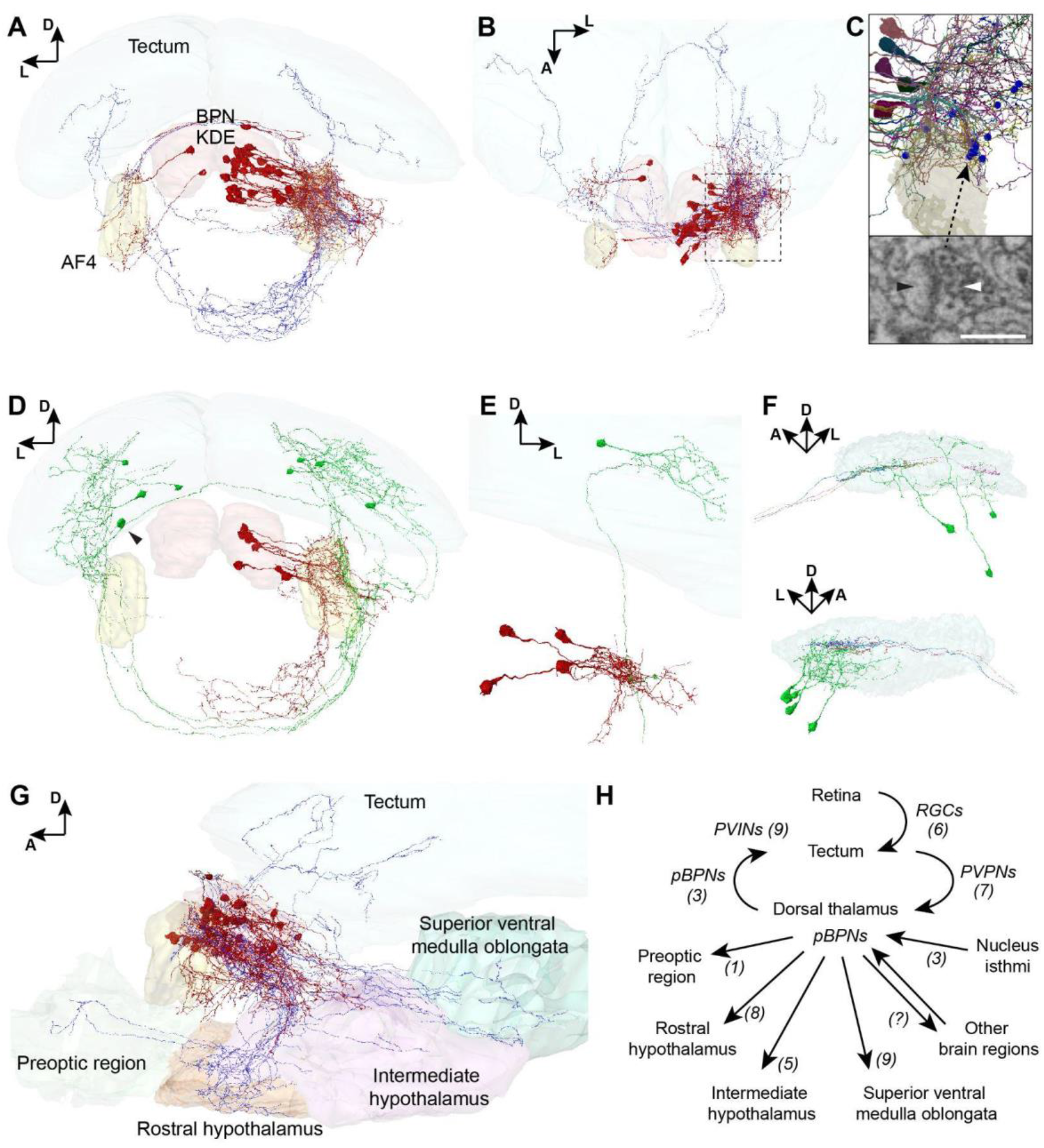
Connectome of the larval thalamic bout-preference region. **a,** Frontal view of an EM reconstruction of neurons in the bout-preference region (BPN KDE, red) of the dorsal thalamus. Axons are shown in blue. **b,** Top view of the neurons shown in **a**. **c,** Close-up of the thalamic arborization field, outlined in **b**. Synapses, of which presynaptic partners have been identified, are shown as blue spheres. One exemplary synapse of a tectal PVPN axon (white arrow) onto a putative BPN’s dendrite (black arrow) is indicated below. Scale bar, 0.5 µm. **d,** Frontal view of tectal PVPNs (green) and their postsynaptic putative BPN partners (red). One pretectal projection neuron is indicated by a black arrow head. **e,** Example of a single PVPN (green), which makes ipsilateral synaptic contacts to at least four identified putative BPNs (red). **f,** Side view of the left (upper panel) and the right (lower panel) tectal SFGS layers, showing the PVPNs (green) and their presynaptic RGC axons (different colors). PVPN axons are not shown for clarity. **g,** Side view of the putative BPNs (red, axons in blue) and their axonal target regions (also see Movie S2). **h,** Circuit diagram. Identified cell types are indicated in italic with cell numbers in brackets.

We further investigated the downstream target regions of putative DT-BPNs. Of our 34 traced neurons, 24 had long projection axons to other brain areas, while 10 neurons had local (n=3) or premature (n=7) axonal projections. Registration of all mapped brain regions from our mapzebrain atlas into the EM dataset revealed the tectum (n=3 cells), contralateral thalamus (n=3), preoptic region (n=1), rostral hypothalamus (n=8), intermediate hypothalamus (n=5), and superior ventral medulla oblongata as axonal targets (n=9; Fig 3g and 3h). Putative BPNs that projected back to the tectum, targeted the SFGS layer, where they contacted tectal periventricular interneurons (PVINs, Fig. S6d).

To complement the EM tracings, we next analyzed the morphology of traced neurons residing in the BPN-KDE of the light microscopic mapzebrain atlas^33^. We identified 13 putative BPNs that all extended their primary neurites ventro-laterally into a neuropil area posterior of AF4, consonant with the EM data. Of these neurons, 12 projected into other brain areas, including tectum (n=1), preoptic area (n=1), rostral hypothalamus (n=1), intermediate hypothalamus (n=3), superior ventral medulla oblongata (n=5), and inferior ventral medulla oblongata (n=2) (Fig. S6e-f).

These findings suggest a pathway for the detection of fish-like motion: Retinal information reaches BPNs via tectal PVPNs and is subsequently transmitted to brain areas proposed to regulate social behavior, including the preoptic region, and clusters in the rostral, intermediate and caudal hypothalamus that showed *c-fos* signal during shoaling behavior (Fig. 1f). Gradual maturation of DT projections and/or addition of synapses, such as those connecting the ventral forebrain at around 14 dpf, may then underlie the emergence of shoaling at the juvenile stage^38^.

### Tectum ablation reduces bout recognition in DT and eliminates attraction

Our inferred wiring diagram places tectal PVPNs upstream of DT-BPNs. Alternatively, DT-BPNs may receive inputs via other sources, such as retinorecipient areas in the thalamus or pretectum^39^. To distinguish between these possibilities, we chemogenetically ablated tectal neurons in larval zebrafish (Fig. 4a) and subsequently recorded calcium responses in the DT-BPN region. We genetically targeted tectal cells for cell ablation using the *SAGFF(lf)81c*^40^ enhancer trap line to drive expression of nitroreductase (*UAS-E1B:NTR-mCherry*^41^). *SAGFF(lf)81c* is strongly expressed in a large fraction of tectal neurons and weakly expressed in parts of the pretectum, habenula, and anterior DT as well as anterior VT (Figs. 4b, S7). The wiring diagram predicts no major role for these additional areas in driving DT responses.

**Fig. 4.**
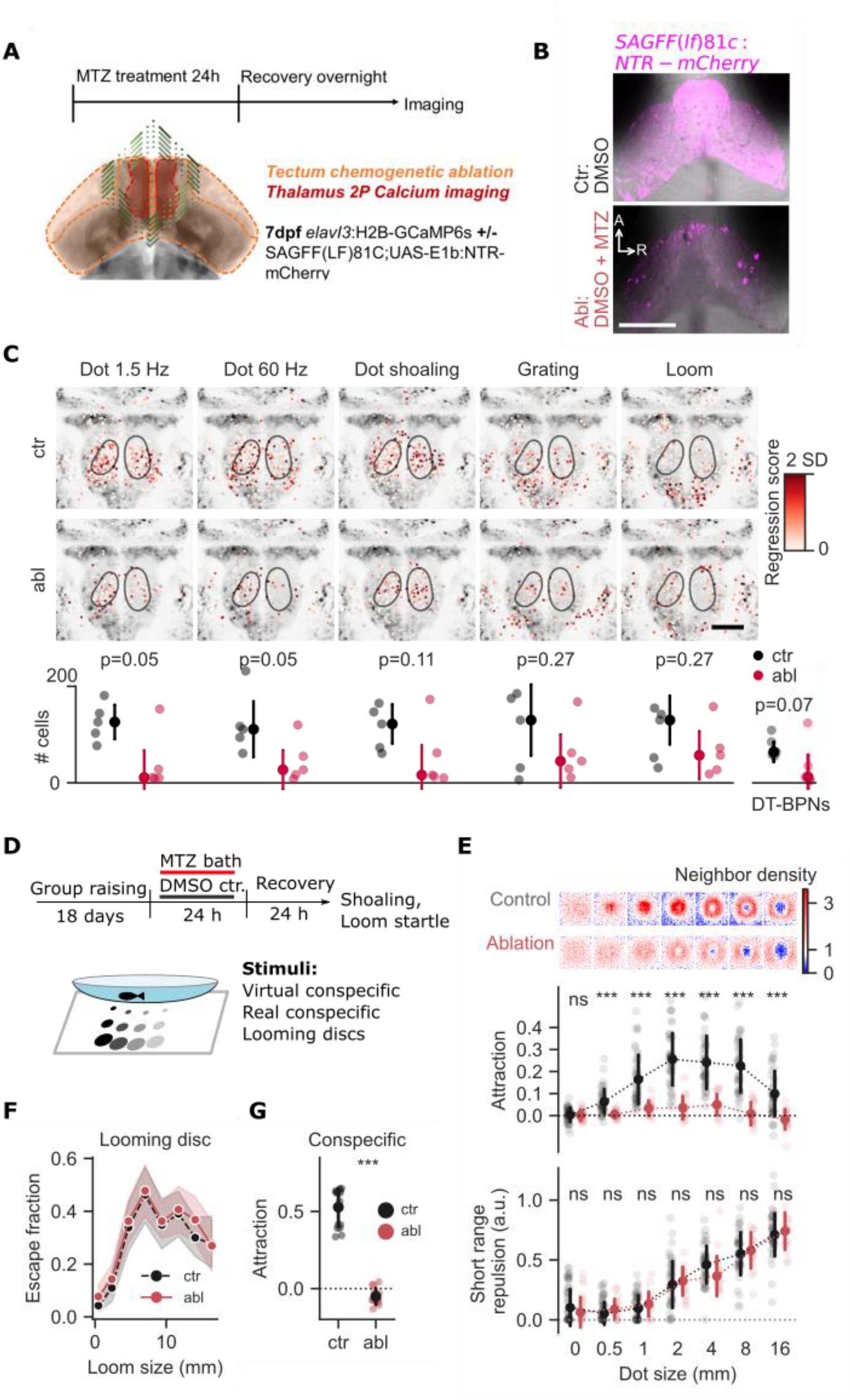
Tectum ablation reduces bout recognition in DT and eliminates attraction. **a,** Schematic of the experimental workflow. 7 dpf larvae were treated with metronidazole (MTZ) for tectum ablation overnight. After 24 hrs recovery, we recorded neuronal responses in DT to bout, continuous and naturalistic motion as well as grating and looming stimuli by volumetric two-photon calcium imaging as in Fig. 2. **b,** Representative epifluorescence images of 7dpf *SAGFF(lf)81c:gal4, UAS:NTR-mCherry* animals 24 hours after treatment with 10 mM MTZ (Ablation) but not DMSO (Control) show extensive cell death throughout the optic tectum. Scale bar: 100 µm. **c,** Tectum ablation reduces bout tuning in DT. Neurons were thresholded based on the 90th percentile of regression scores per stimulus across all animals. Top: Anatomical distribution of top-scoring neurons for dot motion at 1.5 Hz/60 Hz, naturalistic motion (Dot shoaling), grating and looming. Grey background: thalamic region of the mapzebrain reference brain. Outline: BPN KDE, same as in Fig. 2i. Scale bar: 50 μm, n=10 animals. Colorbar: regression score above mean. Bottom: Cell counts of top-scoring neurons per fish for ctr: control and abl: ablated fish. Bottom right: BPN count (BPI>.5) in DT per fish. p-values: one-sided Mann-Whitney U test. Black dots indicate median count ±1SD. **d,** Schematic of shoaling test after ablation of *SAGFF(lf)81c* neurons in 21 day old animals. **e,** Attraction and neighbor density are strongly reduced in ablated animals. Short-range repulsion is intact. N=15 ablated, 45 control animals. Data represent individual animals and mean (attraction) or median (repulsion). Error bars are 1SD. Neighbor maps show mean probability of finding the stimulus in space with the focal animal at the center of the map, heading up. Values represent ratios, relative to time-shuffled data. Each map is 60 x 60 mm. ***: p<0.001, ns: not significant. student’s t-test for attraction, Mann-Whitney U test for repulsion. **f,** Loom induced startle responses are intact in ablated animals. N=14 ablated, 14 control animals. Data represents mean, shading represents 1SD. **g,** Shoaling with real conspecifics is disrupted in ablated animals. Same animals as in **f**. p<0.001:***, student’s t-test.

We found that *SAGFF(lf)81c* ablation reduces the number of DT-BPNs by more than 80% (12±22 vs 64±50 cells per animal) (Fig. 4c). Similar trends are observable for dot motion at 1.5 and 60 Hz, where tectal ablation significantly reduced the number of top-scoring neurons (p=0.05). In contrast, responses to looming and translational grating motion in DT and surrounding pretectum were not significantly affected (p=0.27 and p=0.27), demonstrating specificity of the *SAGFF(lf)81c* ablation.

To test how tectal ablation affects shoaling, we extended the chemogenetic treatment to 21 dpf animals and analyzed free-swimming interactions with virtual conspecifics (Fig. 4d). While ablated animals appeared healthy and had slightly faster swim kinetics, they showed a severe loss of attraction towards virtual conspecifics of various sizes (p<0.001, Fig. 4e, S7b-d). To further investigate the spatial scale of this shoaling defect, we computed neighbor density maps that represent relative spacing with virtual conspecifics. In controls, neighbor maps revealed a central zone of short-range (5-15 mm) repulsion, surrounded by a ring of long-distance attraction (10-30 mm). In ablated animals, this balance was shifted. The ring of attraction was strongly reduced while the zone of repulsion was unaffected (Figs. 4e, S7d,e). Moreover, looming-induced startle responses were intact in ablated animals (p>0.05; Fig. 4f). Finally, we confirmed that tectal ablation disrupted shoaling with a real conspecific (p<0.001; Fig. 4g).

Together, these observations suggest that *SAGFF(lf)81c* neurons are essential elements of a pathway that mediates the affiliative aspect of shoaling, but are dispensable for collision avoidance during shoaling and visual escape from a looming threat^42^.

## Discussion

Affiliation with conspecifics is a core building block of social behaviors that offer collective benefits such as feeding or evasion of predators in swarms^3,8,43^. Consequently, animals need to robustly recognize neighbors to balance attraction and repulsion into an appropriate distance during highly dynamic interactions in cluttered environments^1,4,5^. While empirical models of collective behavior have postulated distinct individual-level behavioral rules^5,7,8^, the neuronal implementation of such coordination was elusive, largely because mutual interactions mask causal relationships between conspecific signals and receiver responses. Our results in shoaling zebrafish highlight fish-like motion^2,11^ as a salient trigger signal of an attraction pathway that converges on a multimodal^35^, socially activated DT cluster and feeds into the conserved social behavior network^12,13^. Neuronal activity in this circuit thus represents an inherently kinetic metric of neighboring animals, unlike current shoaling models that emphasize positional information^1,4,6,23^. In contrast, short-range repulsion engages a separate circuit, likely overlapping with the (non-social) collision avoidance pathway^44–46^. The correspondence of sensory activation in freely shoaling versus immobilized individuals with virtual conspecifics suggests this approach can also reveal the role of individual nodes in the downstream network during shoaling for an understanding how collective dynamics emerge from neuronal computations in individual animals.

## Acknowledgements

We thank Eva Laurell and Enrico Kuehn for generating HCR stainings and for imaging marker lines for the mapzebrain atlas, Joe Donovan for advice in functional imaging and data analysis.

## Author contributions

JMK, KS performed 2 photon imaging experiments. JMK performed registration of 2-photon and HCR data. MJ segmented the EM volume with FFNs. DF proofread and traced presegmented EM data. IS performed HCR staining and imaging. FS generated the EM data and performed EM brain area registration. SS performed pilot HCR *in-situ* experiments in juvenile fish. JL performed behavior experiments, and brain area segmentation. JMK, KS, DF, and JL analyzed the data. JMK, KS, DF, HB, and JL interpreted the data. JMK, DF, HB, and JL wrote the paper with input from all authors. All the authors reviewed and edited the manuscript. HB and JL supervised the project.

## Data and materials availability

All data to evaluate the conclusions in the paper and to reproduce the analysis are present in the paper or made publicly available. Raw HCR data, 2-photon time series for individual neurons and behavior tracking data are available on figshare upon publication. The EM stack will be publicly available upon publication of a companion paper. Python scripts to recapitulate the analysis are available on bitbucket upon publication.

## Supplementary Material

Methods Figs. S1-S7

Movies S1-S3

## Methods

### Animal care and transgenic zebrafish

Adult, juvenile and larval zebrafish (*Danio rerio*) were housed and handled according to standard procedures. All animal experiments were performed under the regulations of the Max Planck Society and the regional government of Upper Bavaria (Regierung von Oberbayern), approved protocols: ROB-55.2Vet-2532.Vet 03-15-16, ROB-55.2Vet-2532.Vet 02-16-31, and ROB55.2Vet-2532.Vet 02-16-122. Incrosses of the following transgenic lines were used: *elavl3:H2B-GCaMP6s* and *SAGFF(lf)81c; UAS-E1B:NTR-mCherry*. Larvae were raised in Danieau solution on a 14/10h light/dark cycle at 28.5 °C until 6 days post-fertilization (dpf). For experiments in juveniles, animals were then raised under standard facility conditions at 28.5 °C in groups of 20-25 individuals. The fish were fed by feeding robots once a day with artemia and 2-3 times a day with dry food.

### Shoaling assay and behavior quantification

Shoaling with real and virtual conspecifics was assayed as previously described^2^. Briefly, 15 or 35 individual animals are transferred individually into shallow watch glass dishes of 10 cm or 7 cm diameter, respectively, separated by a grid of visual barriers and resting on a projection screen. Custom written Bonsai^47^ workflows were used to project stimuli to each animal and to track animal location at 30 frames per second. Stimuli were black dots on white background moving along a predefined, synthetic trefoil shaped trajectory at an average speed of 5 mm/s. For continuous motion, the stimulus position was updated 30 times per second. For bout-like motion, the stimulus position was updated once every 666 milliseconds. Dot size was 4 mm unless noted otherwise, and 0 mm in the no-stimulus condition.

To assay shoaling of pairs of real conspecifics, we introduced a second animal in the same dish and did not show any projected stimuli. FastTrack^48^ was used for post-hoc tracking of real pair shoaling.

Attraction and neighborhood maps were quantified as previously described^2^ using custom written python software: We calculate the ‘real’ average inter animal distance or animal dot distance for each animal in 5 minute chunks (IADr). Next, we generate 10 time shifted trajectories and re-calculate the ‘shifted’ average inter animal or animal dot distance (IADs) for each time shift. Mean IADs for all time shifts is used to compute attraction as (IADs-IADr)/IADs.

For neighborhood maps, neighbor position time series were transformed into the focal animal’s reference frame to compute a binned 2D histogram.

Repulsion was quantified as the reduction in attraction at the center of each animal’s neighbor density map. Neighbor density maps were gaussian filtered (sigma: 3 mm) before obtaining 24 radial line scans (width: 5 mm) starting from the center of the map. Repulsion was the area above the average line scan, at radii less than the radius where maximum neighbor density occurred (Fig. S7e), divided by the full length of the scan (29 mm).

Looming stimuli were presented in the virtual shoaling setup. Looming discs appeared once every minute at a defined offset of 10 mm to the left or the right from the current center of mass of each animal. Looming discs expanded within 500 ms to the indicated final size and did not follow the animal once positioned. To compute an escape fraction we defined an escape response as a trial in which the animal moved more than twice as far in a time window of 1 s immediately following the loom compared to the 1.3 s prior. Bout duration was computed using a custom peak detection algorithm on the velocity time series of each animal.

### *c-fos* activity mapping

#### Shoaling assay for c-fos

For *c-fos* labeling, we used nacre; *elavl3:H2B-GCaMP6s* fish at 21 dpf. 35 fish were transferred into individual dishes and left without stimulation in the presence of white projector illumination from below for acclimatization and to establish a low, non-social *c-fos* baseline. Each animal was assigned randomly to one of the four stimulus groups. After 2 hours, continuous or bout-like motion were shown to groups 1 and 2, respectively, while groups 3 and 4 continued to see no stimulus. After 45 minutes, groups 1, 2, and 3 were quickly euthanized and fixed. Four animals of group 4 were then transferred into the dishes of four other animals of this group for shoaling. After 45 minutes, these eight animals were euthanized and fixed as well.

### HCR staining and imaging

Animals were euthanized and fixed on 4% ice cold paraformaldehyde (PFA). The PFA was washed out after 24 h with 1X PBS and the samples were gradually dehydrated and permeabilized with methanol (MeOH) and stored in -20°C for several days until the hybridization chain reaction (HCR) *in situ* labeling was performed. All the HCR reagents were purchased from Molecular instruments and the staining was performed according to the manufacturer’s protocol for whole-mount zebrafish larvae. Briefly, the samples were separated into 2 juvenile fish per a single 1.5 mL eppendorf tube. Rehydration steps were performed by washing for 5 minutes each in 75% MeOH/PBST (1X PBS + 0.1% Tween 20), 50%MeOH/PBST, 25%MeOH/PBST and finally five times 100% PBST. The samples were permeabilized with 30 µg/mL proteinase K for 45 minutes at room temperature (RT), followed by postfix with 4% PFA for 20 minutes in RT and 5 washes in PBST for 5 minutes each. The samples were pre-hybridized in a 500 µL probe hybridization buffer (Molecular instruments) for 30 minutes at 37°C. Hybridization was performed by adding 2 pmol of each probe set to the hybridization buffer and incubating for 16 hours at 37°C. Probes sets for *c-fos* (*fosab*) (B5 initiator), *cort* (B3 initiator) and *elavl3* (B2 initiator) were purchased from and designed by Molecular Instruments. To remove the excess probes, the samples were then washed 4 times, 15 minutes each, with a wash buffer (Molecular Instruments) at 37°C, followed by 2 washes of 5 minutes each with 5XSSCT (5XSSC + 0.1% Tween20) at RT. Pre-amplification was performed by incubating the samples for 30 minutes in an amplification buffer (Molecular Instruments) at RT. The fluorescently labeled hairpins (B2-488, B3-647, B5-546) were prepared by snap cooling: heating at 95°C for 90 seconds and then cooling to RT for 30 minutes. Hairpin solution was prepared by adding 10 µL of the snapped-cooled hairpins (3 µM stock concentration) to a 500 µL amplification buffer. The pre-amplification buffer was removed, and the samples were incubated in the hairpin solution for 16 hours at RT. The excess hairpins were washed 3 times with 5XSSCT for 20 minutes each wash, and the samples were stored in 5XSSCT in the dark at 4°C until imaging.

For dorsal imaging, the samples were embedded in 2.5% low melting agarose in 1XPBS. Imaging was performed with Leica SP8 confocal microscope equipped with 20X water immersion objective. z-stacks, composing 4 tiles, covering of the entire brain were taken (final stitched image size: 1950 x 1950 px, 1406 x 1406 µm, 3 µm in z). All the 32 samples were imaged with the exact same laser power, gain, zoom, averaging and speed to faithfully quantify and compare the fluorescent signal between the samples. For ventral imaging, the samples were removed from the agarose and dissected in order to remove the jaw and the gills. After the dissections, the samples were embedded upside down and imaged in the same manner. Four brains were lost during ventral imaging and were thus excluded entirely from subsequent analysis.

### Image registration

Image registration was performed using Advanced Normalization Tools (ANTs) (ref) running on the MPCDF Draco Garching computing cluster. Prior to registration, stacks were batch processed in ImageJ. Each stack was downsampled to 512 px width at the original aspect ratio using bilinear interpolation, split into individual channels and saved as .nrrd files. For ventral stacks, artefacts of the dissection such as left-over autofluorescent muscle fibres and skin were masked before registration. Initial attempts to register the *elavl3* HCR channel of dorsal or ventral HCR confocal stacks to a live-imaged two-photon reference of *elavl3-H2B-GCaMP6s* expression were not successful, likely due to deformations resulting from the HCR protocol and diverse qualitative differences in image features between the imaging modalities. Instead, separate dorsal and ventral HCR registration templates were generated from scratch by running *antsMultivariateTemplateConstruction2.sh* on 3 manually selected stacks, respectively. Next, all dorsal and ventral stacks were registered to their respective templates using *antsRegistration*. Finally, the ventral template was registered to the dorsal template using affine + b-spline transformations via *antsLandmarkBasedTransformInitializer* with the help of 25 manually curated landmarks in each stack before applying standard *antsRegistration*. The resulting ventral to dorsal transform was then applied to re-register all ventral stacks into one common (dorsal) reference frame.

### c-fos activity quantification

Image analysis was performed using custom scripts in Python. Registered dorsal and ventral stacks were merged as the arithmetic mean intensity for each animal. To normalize for a drop in signal intensity with tissue depth, the *c-fos* signal was divided voxel-wise by the *elavl3* HCR signal filtered by a 3D gaussian (filter width: 55, 55, 15 µm x,y,z). To identify activity clusters, merged stacks from all animals per condition were generated by finding the maximum intensity at each voxel across animals. A combined RGB hyperstack was generated that showed *c-fos* signal for each condition, *cort* HCR, and *elavl3* HCR for reference in different colours for visual inspection. Activity clusters were manually drawn as 3D masks on the hyperstack using the imageJ segmentation editor on orthogonal overlay views. Masks were drawn with the intent to outline prominent, distinct clusters of *c-fos* signal, irrespective of their modulation by social condition. The full hyperstack, including cluster masks is available (upon publication). Brain areas housing the activity clusters were identified by comparison of the *elavl3* reference to the larval brain atlas mapzebrain^33^ and additional resources^49–51^.

Individual *cort* and *c-fos* positive cells in DT were counted manually using the imageJ cell counter plugin. For statistical analysis across activity clusters and conditions, bulk normalized *c-fos* signal was computed as the average intensity of all voxels belonging to a given cluster. Effect size was determined in each cluster for each condition versus the no-stimulus condition by pair-wise computation of Cohen’s d defined as the difference of the means divided by the pooled standard deviation. To determine significant activity modulation compared to the no-stimulus condition, we performed repeated two-tailed t-tests and corrected for multiple comparisons in each family of tests (each activity cluster) using the Bonferroni correction. Hierarchical clustering of the activity clusters was performed on the effect sizes using the seaborn method ‘clustermap’ with default parameters for average euclidean clustering.

### Functional 2-Photon Calcium Imaging

Two-photon (2P) functional calcium imaging was performed on 6-8 dpf larvae and 17-22 dpf juvenile *elavl3*:*H2B-GCaMP6s* transgenic fish without paralysis. The 6-8 dpf larvae were embedded in 2% agarose with the tail freed. The 17-22 dpf juveniles were embedded in 3% agarose, mouth, gills, and tail were freed using scalpels. Additional oxygen was supplied by continuously perfusing the dish with freshly oxygenated fish water. The embedded fish were mounted on a stage at a custom-built MOM 2P microscope. Only fish that did not drift up or down for a duration of at least 5 recordings were used for analysis. Additionally, fish in which no tectal responses could be observed were eliminated from the analysis. Volumetric imaging of the tectum and/or thalamus was performed using the custom-build MOM 2P microscope with remote focusing, resonance X mirror, galvo Y mirror, 16x objective (Nikon CFI70, NA 0.8, WD 3.0 mm) in the remote arm and 20x objective in the imaging arm (Olympus XLUMPLFLN, NA 1.00, WD Methods 19 2.0 mm). The quick refocusing done by the remote arm enabled rapid sequential imaging of 6 planes with a non-linear step size ranging from 6-24 µm at 5 volumes per second. Remote focusing was left out for the high resolution single plane imaging in Fig. 2 H-I. The plane size ranged between 370×370 µm for larvae to 1075×1075 µm for juveniles. Laser power ranged between 12.3 - 15.4 mW. The spatial sampling (0.7-2.1 µm/pixel) and optical resolution allowed discrimination of single cells with cell body diameters typically in the range of 5-8 µm.

#### z-Stack acquisition and image registration

For each functionally imaged fish, a z-stack of the entire brain was taken (512×512 or 1024×1024 pixels, 2 µm in z, 835-920 nm, plane averaging 50-100x) with the 2P microscope. Larval data was registered to the MapZeBrain atlas^33^ using the *elavl3:H2B-GCaMP6s* reference. For juvenile data, a standard brain was generated from three high-quality z-stacks (150x frame averaging) as described in the *c-fos* section and each juvenile brain was registered two it. The generation of a standard brain and the parameters used for ANTs registration have been described in detail in^33^.

Two align functional ROIs from 2p data to a common reference frame, a 2-step strategy was used. First, average frames (using SDs of pixel intensities) of all imaging planes were generated. Average frames were registered to individual z-stacks by using a template-matching algorithm in a custom written Python script. Then, converted ROI locations in z-stack space were transformed to the larval and juvenile common reference frames by running the ANTs command *antsApplyTransformsToPoints* with the matrices from the z-stack registrations.

#### Visual stimuli

Visual stimuli were designed using PsychoPy projected by a LED projector (Texas Instruments DLP Lightcrafter with 561 nm filter) on Rosco tough rolux diffusive paper placed into a petri dish filled with fish water.

#### Frequency tuning

A black dot moving in a circular trajectory (radius 18 mm) with the fish head in the center was shown starting either perpendicular to the fish at the left, or in the front of the fish. Since no important differences between the starting positions were observed, both were combined in further analysis. The dot was moved in discrete jumps at 0.75, 1.5, 3.0, 6.0 or 60.0 Hz at an overall speed of 5 mm/s. Each frequency was presented using a dot radius of 4 mm. In addition, the behaviorally most attractive (1.5 Hz) and least attractive (60.0 Hz) stimuli were also presented using a dot radius of 2 mm and 8 mm. Both clockwise and counter-clockwise presentations were shown. The frequency, direction and if applicable size were randomly drawn at each stimulus instance. Each stimulus had a duration of 22.6 s and was followed by a 20 s break. A total of 13 stimuli were shown per 10 minute recording. Five to nine of these recordings were performed in each fish, leading to an average of four to six presentations of each stimulus.

#### Specificity

Naturalistic stimulus trajectories consisted of a dot (4 mm radius) moving along real trajectories from one of two interacting juvenile zebrafish, taken from^2^. The trajectory was computed as a fish-centric view of the conspecific with respect to a focal fish. To avoid noise in the heading calculation due to tracking jitter, the trajectory was convolved with a normalized hamming kernel (mode: *valid,* window length: *20*). The naturalistic motion sequences were shown for one minute each. For the whole-field motion stimulus an image was created by combining random intensities and restricted spatial distributions in Fourier space, matching the size of the moving dot. The computed image either rotated in discrete jumps of 1.5 Hz or continuously at 60 Hz (projector frame rate). In both cases the stimulus took 22.6 s to finish a complete round. All stimuli, 1.5 Hz dot, 60 Hz dot, 1.5 Hz whole-field, 60.0 Hz whole-field and naturalistic dot motion were shown in a pseudo-random order during 6×10 min recordings.

#### Kinetic parameters

All shown dots were 4 mm in diameter and moved in the clockwise direction. Five different speeds were tested using a continuously moving dot: 2, 5, 15, 50, and 150 mm/s. Five speeds at a bout frequency of 1.5 Hz were tested by increasing the distance the dot moved during each bout. This increased both the average speed and the acceleration during bouts. The following parameters were tested: 1.25 mm/s; 3 m/s^2^, 2.5 mm/s; 6 m/s^2^, 5 mm/s; 12 m/s^2^, 10 mm/s; 24 m/s^2^, and 20 mm/s; 48 m/s^2^. Finally, for changing acceleration during each bout, we modelled each bout as a gaussian speed profile and changed the width of the curve. Each stimulus still had an average speed of 5 mm/s through a normalization factor. The following peak accelerations were tested: 0.0, 0.02, 0.5, 2.0 and 12.0 m/s^2^.

#### Control stimuli after tectal ablation

Control stimuli consisted of translational gratings moving rostrocaudally with respect to the fish (width: 20 mm, frequency 0.12 Hz, duration 20 sec) and a looming stimulus (expansion from 0.6° to 110° visual angle in 83 ms, delay 10 s with disk and 20 s without stimulus) centered below the fish. One grating was shown at the beginning, followed by the dot stimuli, another grating and finally the looming stimulus. These recording sessions took 10 min each and were separated by a 1 minute break to avoid potential habituation or response suppression due to the looming stimulus.

### Data analysis for 2-photon imaging

Suite2P^52^ was used for motion correction, ROI detection, cell classification and signal extraction. For the entire analysis, the GCaMP6s time-constant of 7 s was used. Based on visual inspection of the raw data, a cell diameter of 4-6 pixels was used. In detail, raw recording files were de-interleaved into separate time series for each plane. An extra motion correction step was required because of ripple noise stemming from the resonance mirror: to avoid alignment to the noise pattern, rigid and non-rigid motion correction was performed on a spatially low-pass filtered time series (Gaussian, sigma=4). The resulting motion correction parameters were applied to the raw data. Next, the time series were down-sampled 5-fold to 1 volume/s. On the downsampled data, ROIs were detected, fluorescent traces were extracted and the ROIs were classified using the custom classifier for *elavl3:H2B-GCaMP6s* neurons.

#### Mean ΔF/F responses

For each functional ROI, the fluorescent trace was normalized and split into stimulus episodes. ΔF/F was computed by using the 5 s prior to stimulus onset as baseline. ΔF/F temporal responses were averaged across stimulus presentations per stimulus and then averaged over time to receive one value per stimulus.

#### Bout preference index (BPI)

Based on the behavioral tuning curves to bout frequency^2^, stimuli were split into bout-like (0.75-3 Hz) and continuous (6-60 Hz) categories. BPI was defined as the difference in mean over mean ΔF/F to bout-like stimuli and mean over mean ΔF/F to continuous stimuli divided by their sum (Equation 1). Bout preference neurons were considered all ROIs that scored BPI>.5, which equates to threefold higher bout response.

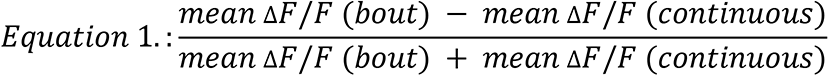

#### Principal Component Analysis

Mean ΔF/F responses to all 18 stimulus conditions from the frequency tuning experiment were taken per neuron to generate a matrix Neurons (=rows) X Responses(=columns). PCA was computed on this matrix using *sklearn.decomposition.PCA*.

#### Tuning peaks

For computing peaks in the tuning of neurons to a variable, mean ΔF/F responses were interpolated with a 1-D spline (*scipy.interpolate.InterpolatedUnivariateSpline*, k=2, second degree) and the location of the maximum was computed.

#### Regression analysis for tectal ablation experiment

To retrieve regression scores for each stimulus from neuron traces, a boxcar regressor was constructed for each stimulus over the whole recording time that was by default 0 and changed to 1 during stimulus presentations. The regressor was then convolved with an exponentially decaying kernel, using the 7 sec time constant of GCaMP6s. To score neurons for each stimulus regressor, a linear regression model was generated (*sklearn.linear_model.LinearRegression*). For the regression score, the regression coefficient was multiplied with coefficient of determination R^2^. To count top-scoring neurons for the control and ablated condition, the overall 90th percentile of all regression scores across experiment conditions was computed separately for each stimulus. Neurons scoring higher than this threshold were counted for both experimental conditions per fish.

#### Gaussian kernel density estimation

To generate a kernel density estimate of BPNs in anatomical space, BPN coordinates were used to fit a Gaussian Kernel (*sklearn.neighbors.KernelDensity*(*\**, *bandwidth=10(*14 for 7dpf), *algorithm=’auto’*, *kernel=’gaussian’*, *metric=’euclidean’*). In detail, the brain was divided along the rostrocaudal axis, and for each hemisphere a separate kernel was fitted with the contained BPNs, using the BPIs as weights. The resulting two kernels were used to generate probability density fields of each hemisphere, which were then merged again. The resulting density was thresholded so that only voxels within the brain itself had values >0 and all voxels in the volume surrounding the brain equaled 0. Probability values were then normalized so that the sum would result in the total number of BPNs. To draw contours of areas with certain threshold BPN density, the KDE volume was binarized so that all voxels above threshold equaled 1. Of the resulting binarized volume a 2-D maximum intensity projection was computed for each orthogonal anatomical axis and a contour-finding algorithm (*skimage*.*measure.find_contours*) was applied to the 2-D projection.

### Definition of the larval DT

The outline of the larval thalamus proper was refined with expert help of Dr. Mario Wullimann, LMU. The refinement was based on extensive analysis of gene expression^49^. The *elavl3* reference stain was used to identify the diencephalic regions. Proliferative cells, however, which are abundant in the anterior DT at the larval stage, are not labeled by *elavl3*. The neurogenin line was used to indicate the early glutamatergic cells belonging to DT. Neurogenin is absent in the prethalamus (VT). The VT/DT boundary was further defined using *gad1b* and *dlx4*, which label late and early GABAergic cells, respectively. GABAergic cells are mainly found in VT, although the intercalated nucleus and the anterior nucleus of DT may contain some *gad1b* positive cells. The pretectum/DT boundary was defined using *gad1b* and *th*. The latter marks dopamine cells present in the pretectum.

### Electron microscopy and segmentation of mapzebrain regions

A detailed description of the EM dataset and region mapping will be published elsewhere (Svara et al., in preparation). Briefly, the Serial Block Face Scanning EM dataset was of a 5 dpf larval zebrafish imaged at a resolution of 14 x 14 x 25 nm. A diffeomorphic mapping between the mapzebrain light-microscopy brain reference coordinate system and the EM coordinate system generated by the dipy (dipy.org) python library was used to overlay mapzebrain (http://fishatlas.neuro.mpg.de/) region annotations over the EM data. Registration accuracy was reviewed for different brain regions (see Figure S5) with an alignment error of maximal ∼5 µm (midbrain) to ∼20 µm (hindbrain). We applied flood-filling networks for an automated reconstruction of all neurons^53^ within the whole-brain EM dataset (Svara et al., in preparation). To correct for split and merge errors of the segmentation, we used the Knossos application (www.knossos.app). Proof-reading of single pBPN-DT cells started at the cell body location and ended, when all branches were completely traced. Growth cones defined premature neurons. Proof-reading of partner cells started at the synapse and was again performed until the whole cell was completed.

### Nitroreductase ablations

To chemogentically ablate neurons, fish were treated with 7.5 mM Metronidazole (MTZ) in Danieau’s solution or fish facility water for larvae and juvenile fish, respectively, with 1 mL/L Dimethylsulfoxid (DMSO). Fish with the transgene *UAS:NTR-mCherry* were incubated in the same tank with control fish lacking the transgene for at least 16 hours overnight. Animals recovered for 24 hrs before starting either imaging or behavior experiments.

### Statistical analysis

All analyses were performed with custom-written Python code, using NumPy, Scipy, MatplotLib, Suite2p, Pandas, Scikit-learn. All statistical details are described in the figure legends and the material and methods. All tests were two-tailed, unless noted otherwise. Error bars represent standard deviations, unless noted otherwise. N denotes number of animals, unless noted otherwise.

---

**Fig. S1.**
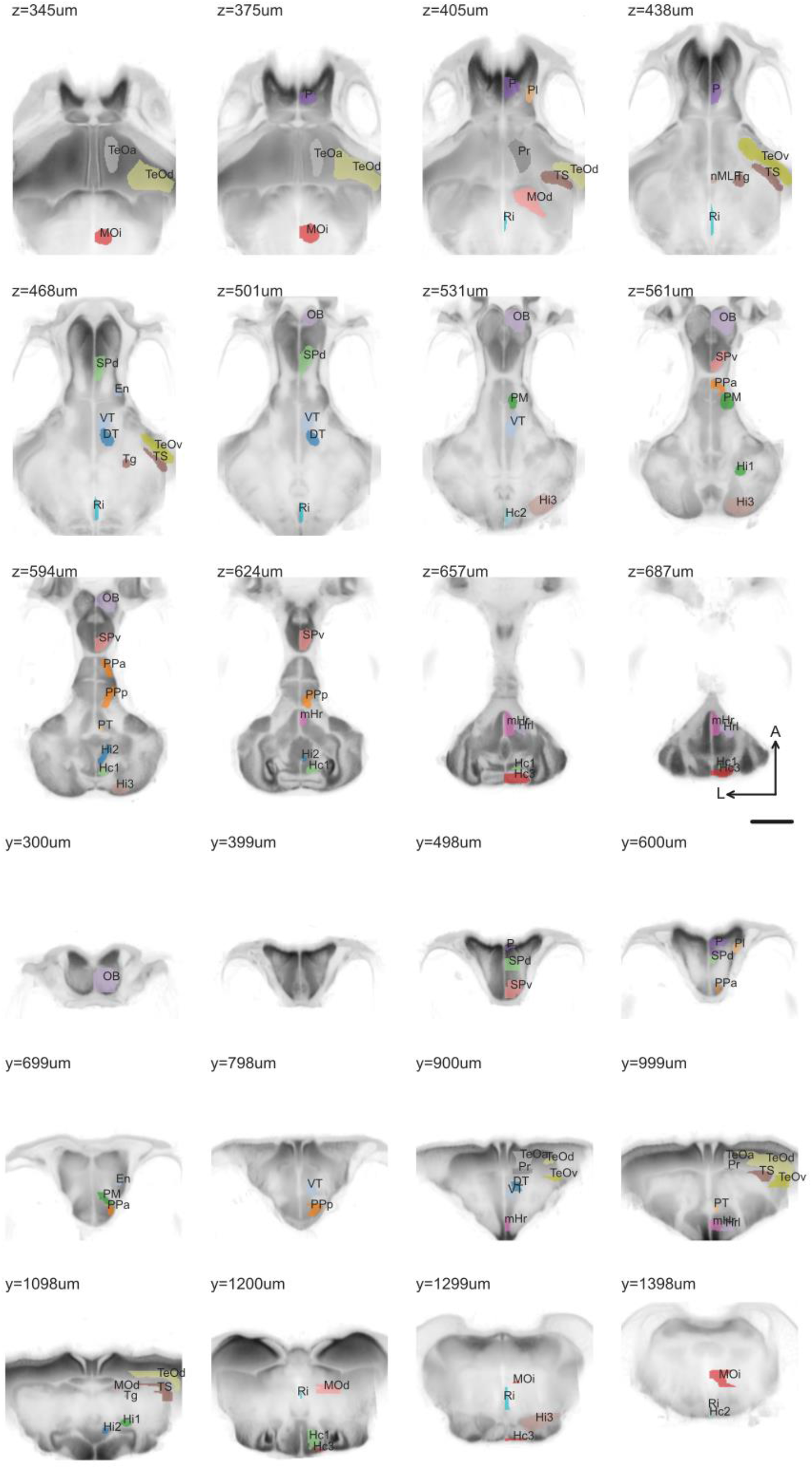
Horizontal and coronal slices showing *c-fos* activity clusters overlaid on the average registered *elavl3* signal from all 28 animals. For visualization purposes, the *elavl3* signal was non-linearly transformed using a gamma adjustment of 0.5. Top three rows are horizontal sections, bottom three rows are coronal sections. Scale bar: 200 μm.

**Fig. S2.**
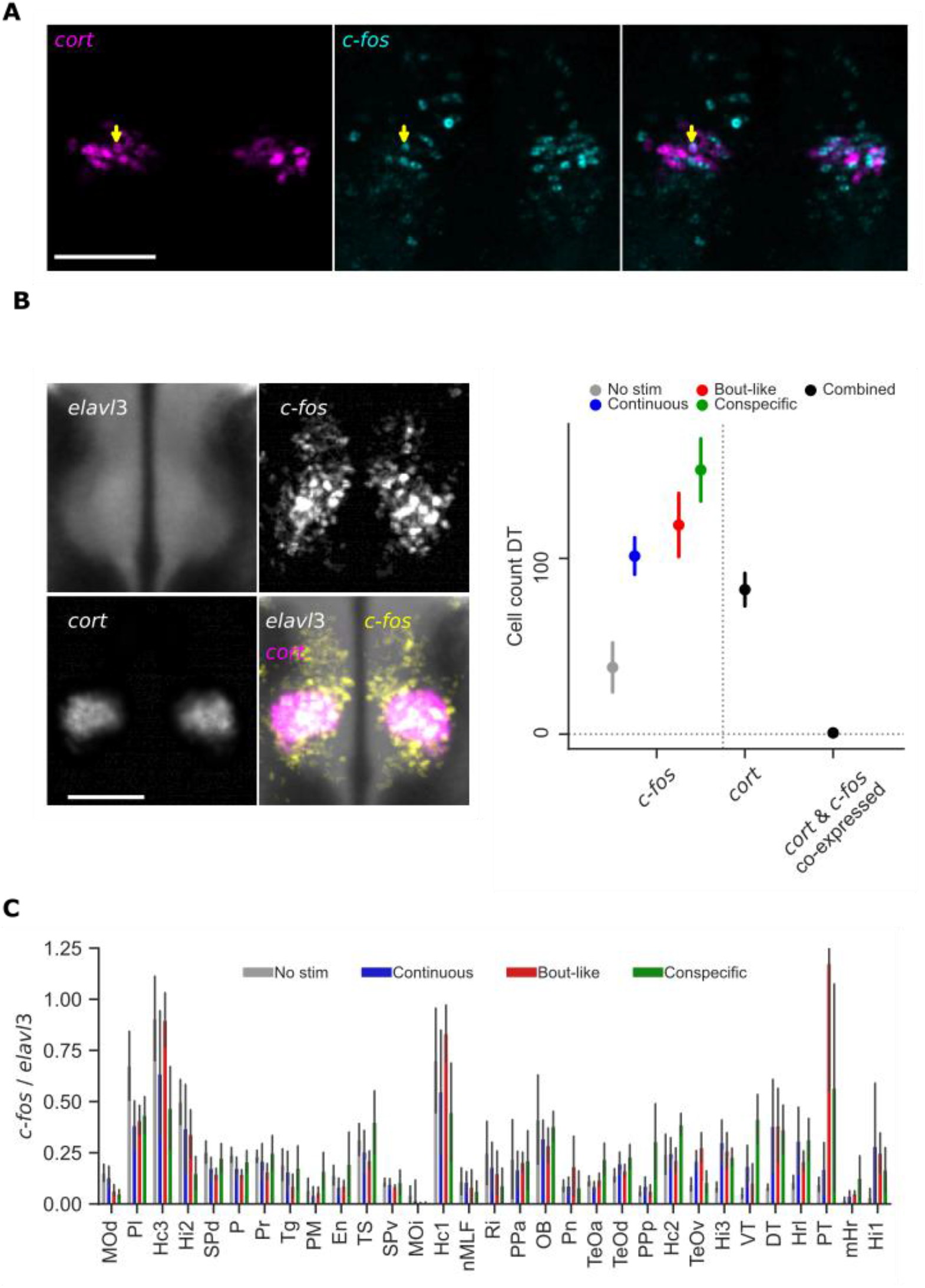
**a,** Representative dorsal thalamic plane showing HCR labelling of *c-fos* and *cort* in one animal after interaction with a bout-like motion dot stimulus. Expression occurs in the same regional cluster (DT) but only one cell expresses both markers (arrowhead). Scale bar: 100 μm. **b,** Left: Expression of *cort* and *c-fos* induction by shoaling stimuli localize to the same area. *elavl3*, *cort* and *c-fos* were co-labeled in the same animals. A single horizontal imaging plane at the center of the DT cluster is shown. *elavl3* and *cort* channels are mean intensity, *c-fos* channel is maximum intensity over all 28 animals. Scale bar: 100 μm. Right: *c-fos^+^* and *cort*^+^ cells were counted in the dorsal thalamus of n=6 animals per condition. Data are mean±1SD. **c,** Normalized bulk *c-fos* signal for each activity cluster. Bars represent mean±1SD, n=6-8 animals per condition.

**Fig. S3.**
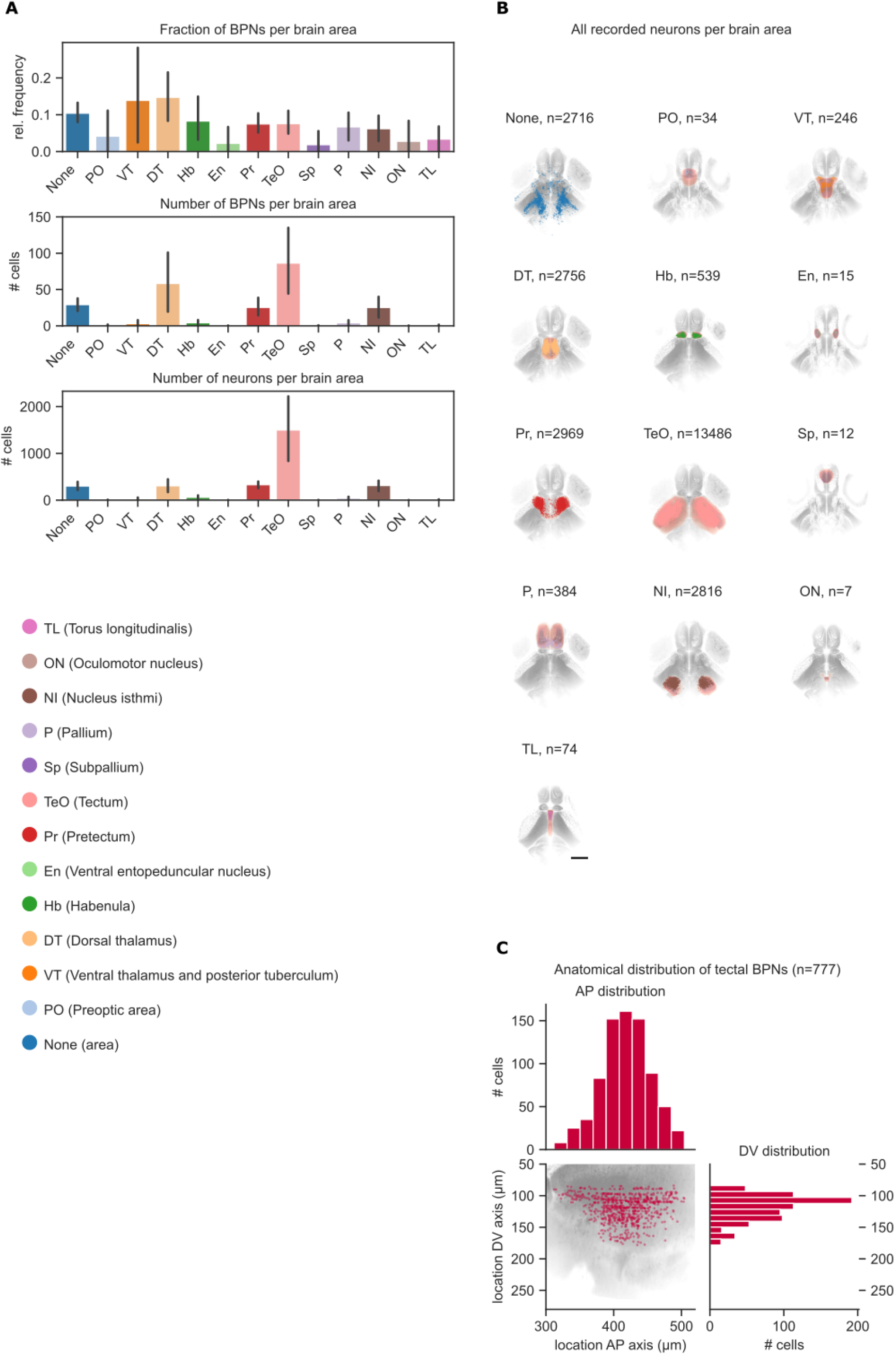
**a**, Top: Relative frequency of BPNs per brain area. Middle: Total number of BPNs per brain area. Bottom: Total number of recorded neurons per brain area. N=9 animals, error bars are 1SD. Scale bar: 200 μm. **b,** Distribution of recorded neurons in each brain area. Red masks are brain areas, dots represent individual neurons. **c,** Distribution of tectal BPNs along anterior-posterior (AP) and dorso-ventral (DV) axis.

**Fig. S4.**
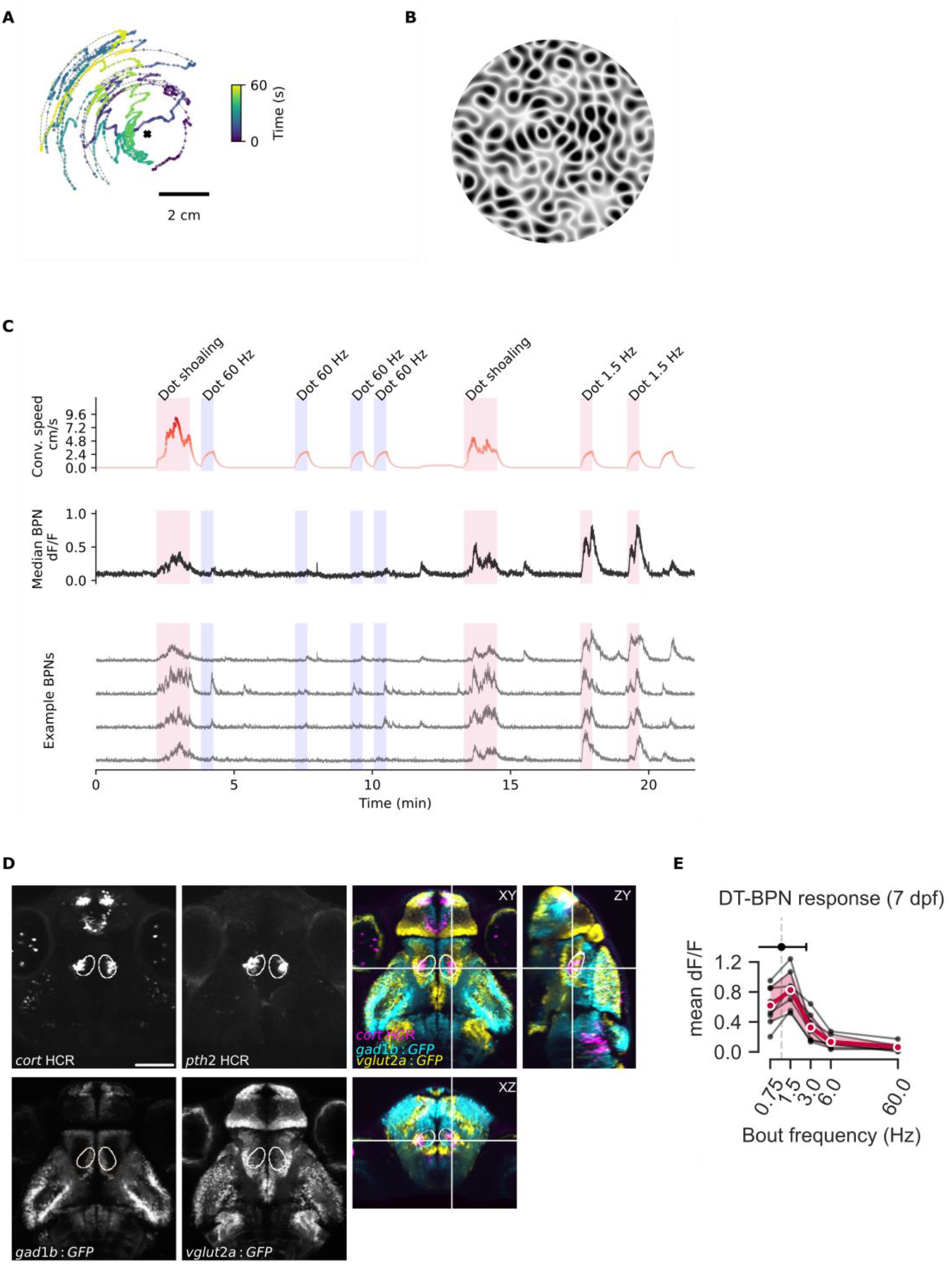
**a**, Dot shoaling stimulus. Dot position recapitulates the location of a conspecific relative to a focal fish facing up. Cross marks location of the focal fish. **b,** Whole-field motion stimulus. **c,** DT-BPNs respond to dot shoaling stimulus. Red trace shows instantaneous dot stimulus speed convolved with GCaMP6s kernel (see methods). Dot shoaling, 1.5 Hz and 60 Hz stimuli were shown in pseudo-random order as indicated. Median trace represents n=72 BPNs from one fish, including four representative neurons shown below. **d,** Expression of *cort*, *pth2*, *gad1b* and *vglut2a* defines the location of the larval BPN KDE as dorsal thalamus. *gad1b* positive ‘stripe’ of cells near the midline marks the dorsal edge of VT. Left four panels show single planes. Right shows merged orthogonal views. Each channel shows mean expression over 3-5 individual fish registered to the mapzebrain atlas. *Cort* and *pth2* are HCR labels. *Gad1b* and *vglut2a* are gal4 enhancer traps lines driving expression of GFP. Scale bar: 100 µm. Also see Movie S1. **e,** Larval mean DT-BPN (n=40±23 per fish) tuning curve to stimulus frequencies from 0.75 to 60 Hz shown in red. N=8 animals. Mean tuning peak of individual neurons (shown above) was 1.1 Hz±1.5 Hz, matching juvenile zebrafish bout frequency (n=326 neurons). Black lines represent means of individual animals. Data from subset of all animals in Fig. 2J with number of recorded DT-BPNs>10.

**Fig. S5.**
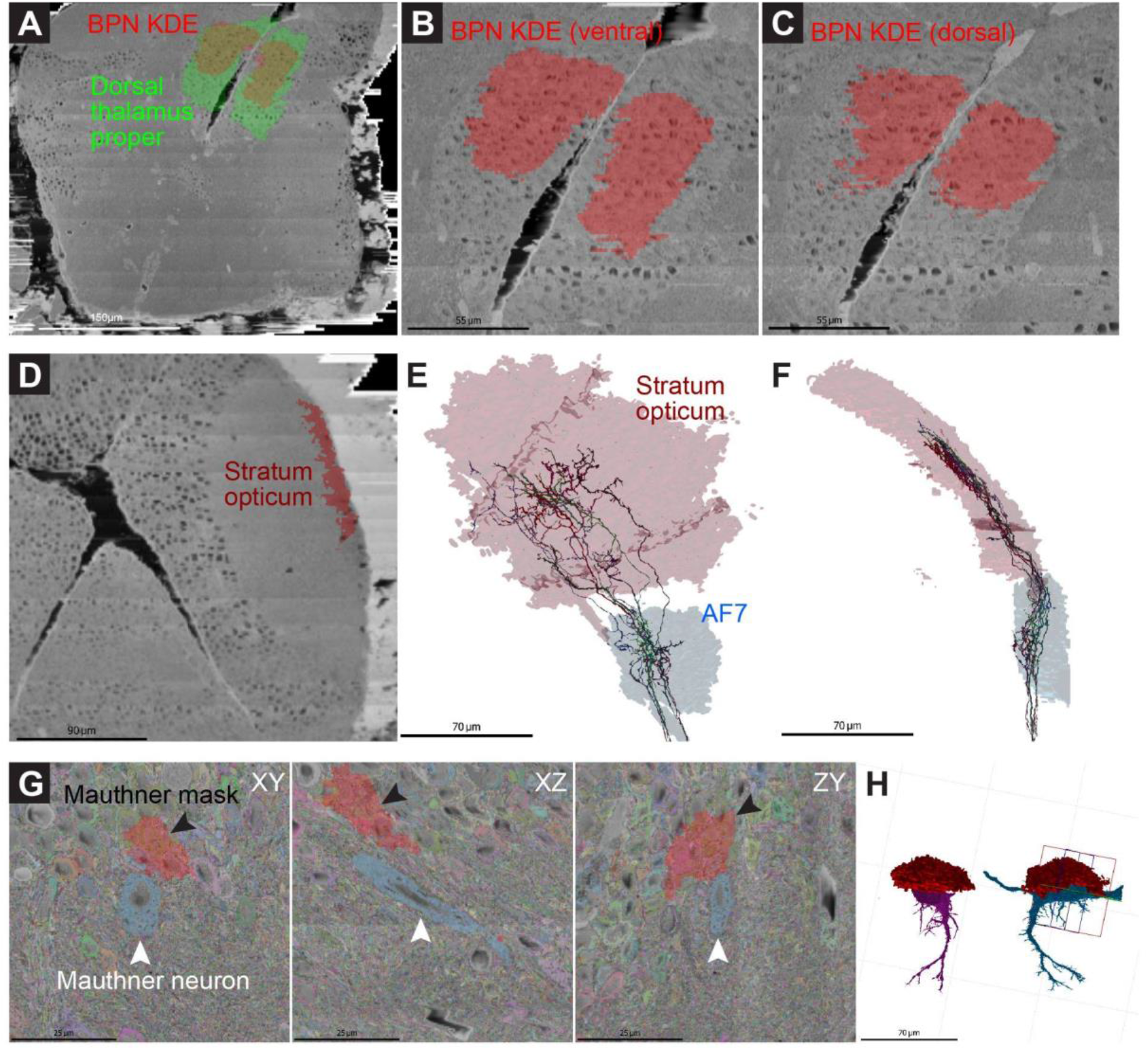
Registration of mapzebrain regions to the EM dataset. **a-c:** Top-views of the BPN KDE (red) inside the dorsal thalamus proper (green). **d-f:** Additional examples for the accuracy of registration. Retinal ganglion cell axons, which have been traced in the EM dataset and which project both to the tectal stratum opticum (SO) and the arborization field 7 (AF7), localize exclusively inside the registered regions. **g-h**: The registered mask for the Mauthner neurons, which was generated in the mapzebrain atlas (red, black arrowheads), deviates by ∼20 µm from the location of the Mauthner cells, recognizable in the EM dataset (blue, white arrowheads).

**Fig. S6.**
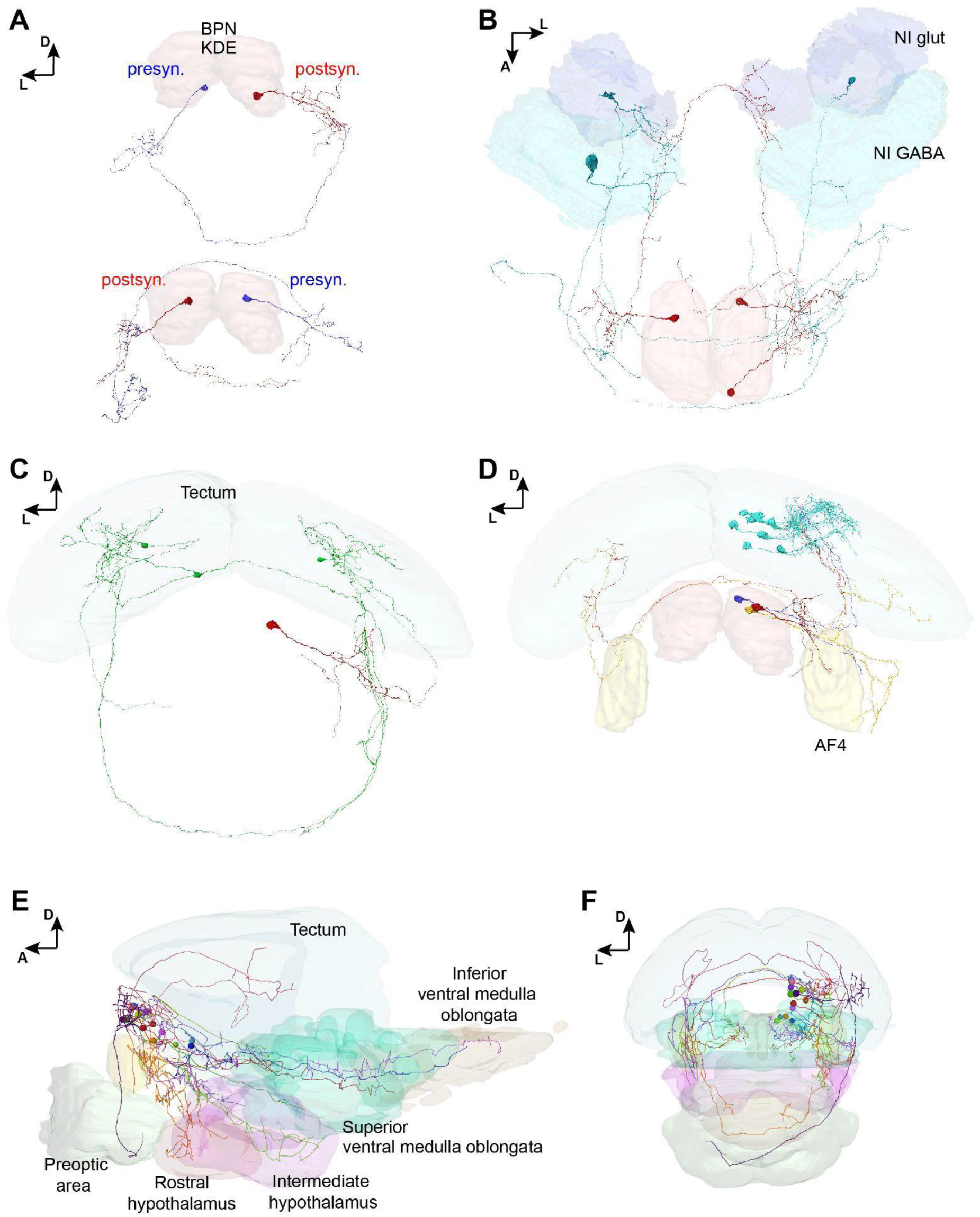
**a,** Frontal views of the BPN KDE in EM reconstructions showing two examples (top and bottom) for synaptically connected putative BPN partners across brain hemispheres. **b,** Dorsal view of three putative BPNs (red) and their presynaptic partners in the nucleus isthmi (glutamatergic and GABAergic domains are annotated). **c,** A single putative BPN (red) receives ipsi- and contralateral synaptic input from at least three identified tectal PVPNs (green). **d,** Three examples for putative BPNs (red, orange, purple) with axonal projections to the ipsi- and contralateral tectum. Eight tectal PVINs (cyan), which are postsynaptic to the red putative BPN are shown. **e-f,** Mapzebrain atlas showing single cells in the BPN KDE and their targeted brain regions in lateral (A) and frontal (B) views.

**Fig. S7.**
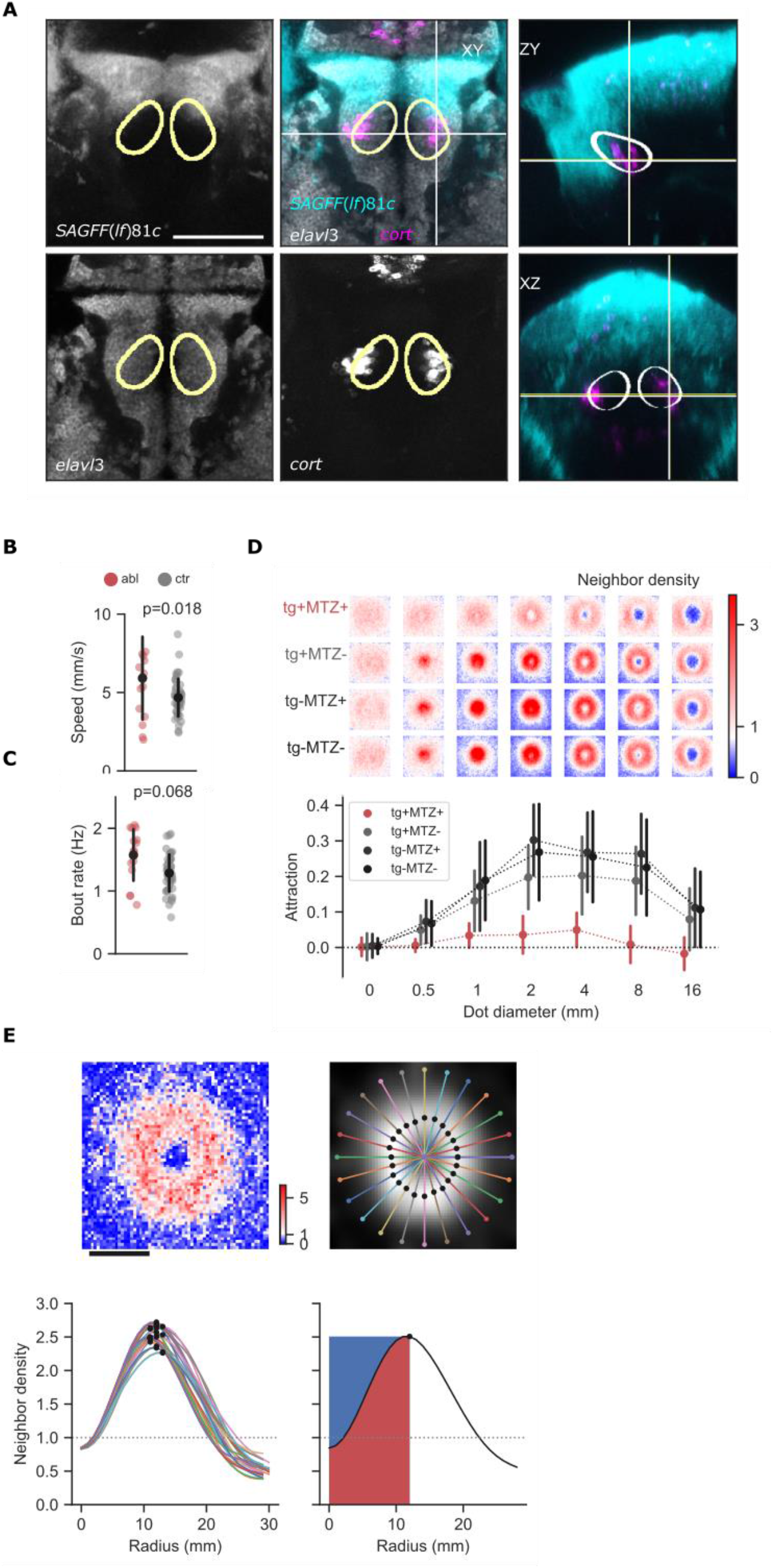
**a,** Expression of *SAGFF(lf)81c:gal4*, *UAS:NTR-mCherry* relative to BPN KDE (yellow outline) and *cort*. Markers were imaged in separate 5-7 dpf fish and registered to the mapzebrain standard brain. *81c*: average of 4 animals *SAGFF(lf)81c:gal4, UAS:NTR-mCherry*. *elavl3*: mapzebrain reference channel. *Cort*: average of 3 animals, HCR label. Merged view is shown in 3 orthogonal planes. White crosshairs indicate the orthogonal planes. Scale bar: 100 µm. Also see Movie S3 **b,** Average swim speed, same animals as in Fig. 4g. **c,** Average bout rate, same animals as in Fig. 4g. **d,** Neighbor density and attraction to dot stimuli as in Fig. 4g, showing each control individually: tg+ and tg- indicate presence and absence of transgene *SAGFF(lf)81c:gal4, UAS:NTR-mCherry,* respectively. MTZ +/- indicates presence/absence of MTZ. Data are mean±1SD. N=15 animals per group. **e,** Definition of short-range repulsion. Top left shows a representative neighbor map for one animal interacting with a 4 mm dot. Scale bar: 20 mm. Color map as in Fig. 4e. Top right and bottom left show 24 radial line scans of smoothed neighbor density. Each scan begins at the center of the map. Black dots label the maximum of each scan. Bottom right shows the mean of all line scans in black. Repulsion is quantified as the area above the mean line, left of the peak (blue shading). This defines the reduction of attraction at the center of the map. Dotted line in bottom panels indicates baseline neighbor density (random distribution).

## Supplementary Movies

Movie S1

Full stack of mapzebrain data shown in Fig. S4d.

Movie S2

EM reconstruction of putative DT-BPNs (red, axons in blue) and their target brain regions. See Figure 3g for region annotations.

Movie S3

Full stack of mapzebrain data shown in Fig. S7a.

